# Electrically induced bacterial membrane potential dynamics correspond to cellular proliferation capacity

**DOI:** 10.1101/542746

**Authors:** James P Stratford, Conor L A Edwards, Manjari J Ghanshyam, Dmitry Malyshev, Marco A Delise, Yoshikatsu Hayashi, Munehiro Asally

## Abstract

Membrane-potential dynamics mediate bacterial electrical signaling at both intra- and inter- cellular levels. Membrane potential is also central to cellular proliferation. It is unclear whether the cellular response to external electrical stimuli is influenced by the cell’s proliferative capacity. A new strategy enabling electrical stimulation of bacteria with simultaneous monitoring of single-cell membrane potential dynamics would allow bridging this knowledge gap and further extend electrophysiological studies into the field of microbiology. Here we report that an identical electrical stimulus can cause opposite polarization dynamics depending on cellular proliferation capacity. This was demonstrated using two model organisms, namely *B. subtilis* and *E. coli*, and by developing an apparatus enabling exogenous electrical stimulation and single-cell time-lapse microscopy. Using this bespoke apparatus, we show that a 2.5 sec electrical stimulation causes hyperpolarization in unperturbed cells. Measurements of intracellular K^+^ and the deletion of the K^+^ channel suggested that the hyperpolarization response is caused by the K^+^ efflux through the channel. When cells are pre-exposed to UV-violet light, the same electrical stimulation depolarizes cells instead of causing hyperpolarization. A mathematical model extended from the FitzHugh-Nagumo neuron model suggested that the opposite response dynamics are due to the shift in resting membrane potential. As predicted by the model, electrical stimulation only induced depolarization when cells are treated with antibiotics, protonophore or alcohol. Therefore, electrically induced membrane potential dynamics offer a novel and reliable approach for rapid detection of proliferative bacteria and determination of their sensitivity to antimicrobial agents at the single-cell level.

## Introduction

Compared to animal bioelectrical signaling, bacterial electrical signaling is understudied and only recently were the excitation dynamics of membrane potential shown to mediate the intra- and inter-cellular signaling which regulates important physiological processes, namely mechanosensation, spore formation and biofilm dynamics (1–4). In animal bioelectrical signaling, externally applied electrical stimuli and measurements of cellular electrical properties have been the principle methodology (5–8). This approach has led to many key discoveries regarding the roles of animal bioelectrical signaling (e.g. early tissue development (9,10), regeneration (11) and carcinogenesis (12–14)) and has fostered the development of real-world applications such as for tissue engineering (15–17), wound healing (6,18) and electroceuticals (19). Utilizing exogenous stimuli is an important step forward toward understanding bacterial electrical signaling and development of applications based on bacterial electrophysiology. In the past, applications of electric currents to bacteria were used for sanitization (20), electroporation (21) and most recently synthetic biology (22). However, due to the only recent discovery of bacterial membrane-potential excitation dynamics, use of external electrical signals in the context of bacterial electrophysiology has been left largely unexplored.

An external electrical stimulus alters cellular membrane potential according to the Schwan equation:

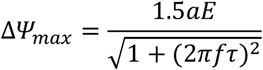

where Δ*Ψ*_*max*_ is the induced membrane potential, *a* is the cell radius, *E* is the applied field strength, *f* is the AC field frequency and *τ* is the relaxation time of the membrane (23). This equation, derived from the electromagnetic theory (24), expresses that the maximum change in the membrane potential of a cell caused by an electrical stimulus is proportional to the applied field strength. Theoretically, when an electrical stimulus is applied to proliferative bacterial cells, it should lead to opening of voltage-gated K^+^ channels (Kv) and consequent hyperpolarization due to K^+^ efflux. Conferring the typical values of bacterial resting potential (−140∼−75 mV (25,26)) and threshold potential for Kv (∼−50 mV (27)) on the equation, one can expect that the depolarization by an electrical stimulus with the field strength of approximately +35∼120 mV/μ m should open voltage-gated K^+^ channels on bacteria.

In addition to its role in bioelectrical signaling, membrane potential is central to cellular proliferation; it provides the essential driving force for ATP synthesis (28) and is crucial for cell division (29). A quantitative estimation based on the measured energy consumption of *E. coli* suggests that the maintenance of membrane potential accounts about half of the total energy consumption (30), and thus it is inherently linked to the proliferative capacity. In fact, the proliferative capacity is commonly determined either by direct time-lapse observation of individual cells or by probing the intracellular state using membrane potential. The latter could be achieved using fluorescent indicators for membrane potential such as Thioflavin T (ThT), DiOC_2_(3) and Rhodamine 123 (3, 31–34). However, the use of such indicators for determining the proliferative capacity is known to be difficult because membrane potential can be affected by many physiological states and environmental conditions (35–37). Due to the baseline fluorescence of cells being affected by a variety of conditions, comparisons between individual cells, populations of cells and difference species are technically complex. This means that meticulous and tedious calibrations are required for species, strains, mediums and detection systems. The difficulties associated with these calibrations often preclude the broad use of these agents for the detection of proliferative bacteria. Nevertheless, these fluorescent indicators provide a useful indication of intracellular physiological state for live-cell imaging.

The dual roles of membrane potential in both signaling and proliferation beg the question regarding the interplay between these two roles of membrane potential. In particular, could the electrical response of cells be affected by the proliferative capacity of individual cells? This is a fundamentally important question for understanding bacterial electrical signaling because the input-output relations are fundamental to “signaling” (38). However, it remains unclear whether cellular responses to an electrical stimulus differ depending on their proliferative capacity. If it does, one may expect that an identical signal input produces different outputs depending on their proliferative capacities, which could be a novel dynamics-based approach to probing cellular proliferative capacity through the monitoring of cellular electrical responses.

In this study, we utilized an exogenous electrical stimulus to investigate the impact of proliferative capacity on electrical signal response. By experimentally testing a prediction from a mathematical model, we showed that an exogenous electrical stimulus induces hyperpolarization in unperturbed cells while inducing depolarization in inhibited cells. This finding offers a novel application to use bacterial electrophysiological dynamics for rapid detection of proliferative cells and differentiation of proliferative and non-proliferative cells within a minute after electrical stimulation.

## Results

### Development of an apparatus that enables the monitoring of membrane-potential response to an exogeneous electrical stimulus

To investigate the potential impacts of proliferative capacity on electrical signaling response, direct observation of cell proliferation, membrane potential and its response to electrical stimuli at the individual-cell level is needed. However, the commercially available apparatus for neural electrophysiology were unsuitable due to the small size of bacterial cells; i.e. ∼1.4 μ m^3^ for *B. subtilis* and *E. coli* (39). To overcome this technical challenge, we designed and developed a novel tool for bacterial electrophysiology. The tool consists of an electrical relay circuit with an open-source I/O board, Arduino UNO, and a bespoke electrode-coated glass-bottom dish (Fig. 1A and S1-3, see Methods for details). Bacterial cells were inoculated on agarose pads and placed on the electrode surface. Importantly, this setup enables the monitoring of cells at single-cell resolution using phase-contrast and the fluorescence membrane-potential indicator, ThT (Fig. 1B).

**Figure 1.**
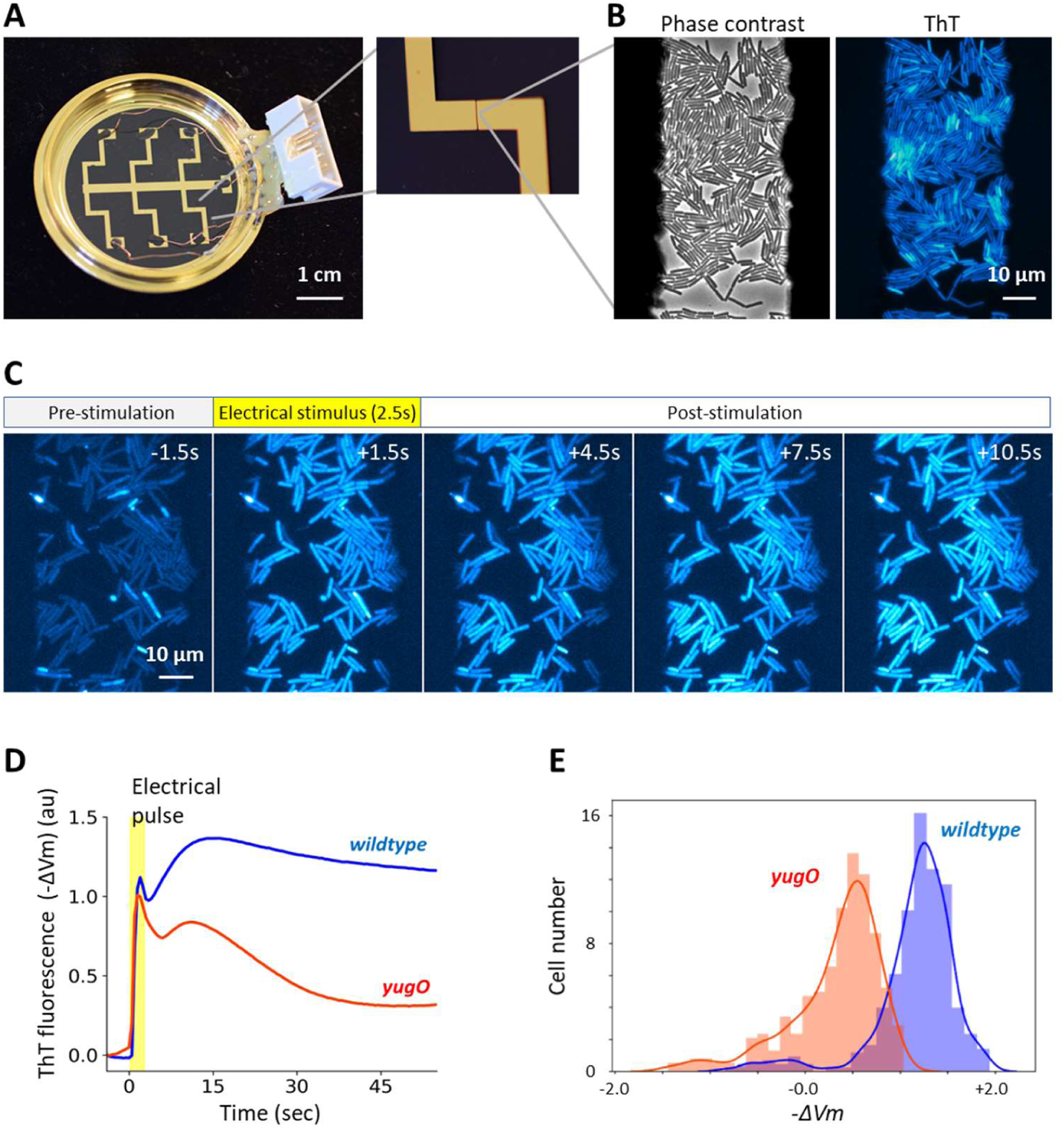
An apparatus enabling concurrent single-cell microscopy and stimulation with exogeneous electrical signal revealed hyperpolarization response to an electrical stimulus. A) Bespoke glass-bottom dish coated with gold-titanium electrodes. Zoomed image on the right shows 50 μ m gap between electrodes. Dish is connected to relay circuit to apply electrical stimulation to bacterial cells (see Fig. S1-3 for details). B) *B. subtilis* cells witnin the 50 μ m electrode gap are visible in phase-contrast and ThT fluorescence images. C) Film-strip images of ThT fluorescence of *B. subtilis* before, during and after electrical stimulation. Increase in ThT fluorescence indicates hyperpolarization response to an electrical stimulus. D) Mean −ΔVm over time for *B. subtilis* wildtype and *yugO* strains. −ΔVm was calculated by log(*F*_*ThT*_*/F*_*ThT,R*_), where *F*_*ThT*_ is ThT fluorescence and *F*_*ThT,R*_ is ThT fluorescence at resting state (see SI text). Time-traces of individual cells are shown in Fig. S4 (wt: n = 321; *yugO*: n =308). Images were taken at 2 fps. E) Histogram of −ΔVm at 30 sec after electrical stimulation. The distributions of wildtype and *yugO* are clearly distinguishable.

### Electrical stimulation causes hyperpolarization of cells via K^+^ efflux

To examine whether an externally applied electrical stimulus is capable of opening K^+^ channels on bacterial membranes, we applied an exogenous electrical stimulus (3 Vpp AC 0.1 kHz for 2.5 sec) to *B. subtilis* cells placed between the 50 μ m electrode gap (Fig. 1B). Upon electrical stimulation, the intensity of ThT fluorescence increased, indicating a hyperpolarization response (Fig. 1C). Single-cell analysis of the fluorescence dynamics revealed that most cells exhibited the hyperpolarization of membrane potential (V_m_), while a small subpopulation of cells depolarizes upon stimulation (Fig. S4A). Intriguingly, the growth rate of these depolarizing cells was found to be much lower compared to other cells (Fig. S5). No detectable change in ThT intensity was observed with the absence of electrical stimulus, indicating that the observe dynamics are induced by the electrical stimulus (Fig. S6). The hyperpolarization response suggests that electrical stimulation causes the efflux of cations such as K^+^, the dominant intracellular cation. It has been shown that chemical depolarization opens the YugO potassium channels (3). The Schwan equation predicts that external electrical stimuli can depolarize bacterial cellular membranes (40). Therefore, we hypothesized that depolarization by the electrical stimulation opens YugO channels, resulting in membrane hyperpolarization by K^+^ efflux.

To test this hypothesis, the same experiment was conducted with a mutant strain lacking the gene encoding the YugO potassium channel. Although the strain still showed an initial hyperpolarization response in the timescale of ∼5 sec, the hyperpolarization response on the timescale of ∼30 sec is greatly attenuated (Fig. 1D, E and S4B). For a further test, we measure the intracellular K^+^ levels using Asante Potassium Green-2 AM (APG-2 AM) (3), and found that the intracellular K^+^ decreases upon stimulation (Fig. S7). These results suggest that K^+^ efflux through the YugO channel is responsible for the hyperpolarization following an electrical stimulation. It also suggests that there may be other voltage-gated channels with faster timescales of activation and inactivation (∼5 sec). It is worth noting that bacteria have several voltage-gated ion channels (1,41,42). This is an interesting observation in conjunction with the fact that different neural ion channels have their unique timescales of activation and inactivation which contribute to information processing (27). We also conducted the same experiment with *E. coli* and confirmed that *E. coli* cells also exhibit hyperpolarization in response to an external electrical stimulus (Fig. S8). Together, these results demonstrate with single-cell resolution that a pulsed electrical stimulus can induce a hyperpolarization response in bacterial cells.

### Exposure to UV-violet light abolishes hyperpolarization response to an electrical stimulus

Having tested our apparatus, we examined the impact of proliferative capacity on signal response by using inhibited cells. To inhibit the proliferative capacity of cells, we chose UV-Violet light (400 nm) because it is one of the most commonly used sanitization methods, which has been shown to be effective with both Gram-positive and Gram-negative bacteria (43,44). Importantly, application of UV-V light allows spatially precise inhibition, creating both irradiated and unirradiated regions within the same field of view. This is critical because it ensures identical electrical stimulation is applied to both proliferative and inhibited cells. We irradiated *B. subtilis* cells in a defined region by UV-V light for 30 sec (Fig. 2A). The growth suppression of the irradiated cells was confirmed by the single-cell analysis of phase-contrast time-lapse microscopy before being stimulated with an electrical pulse (Fig. S10). Upon an electrical stimulation, the irradiated cells exhibit depolarization, while cells in untreated regions become hyperpolarized, in spite of the fact that both received identical electrical stimulus (Fig. 2B and C). This experiment demonstrates that an identical electrical stimulus can result in cellular response in apparent opposite directions depending on whether cells are exposed to UV-V or not. Strikingly, analysis of the fluorescence dynamics after electrical stimulation showed a clear bimodal distribution correlating with the irradiation (Fig. S11A and B). To examine whether this is unique to *B. subtilis*, we conducted the same experiment with *E. coli* cells. The result with *E. coli* also revealed distinct responses depending on whether cells were treated by UV or not (Fig. S11C). These results suggest that proliferative and growth-inhibited cells respond differently to an identical electrical stimulus and that this difference in response dynamics is common to these two phylogenetically distant model organisms.

**Figure 2.**
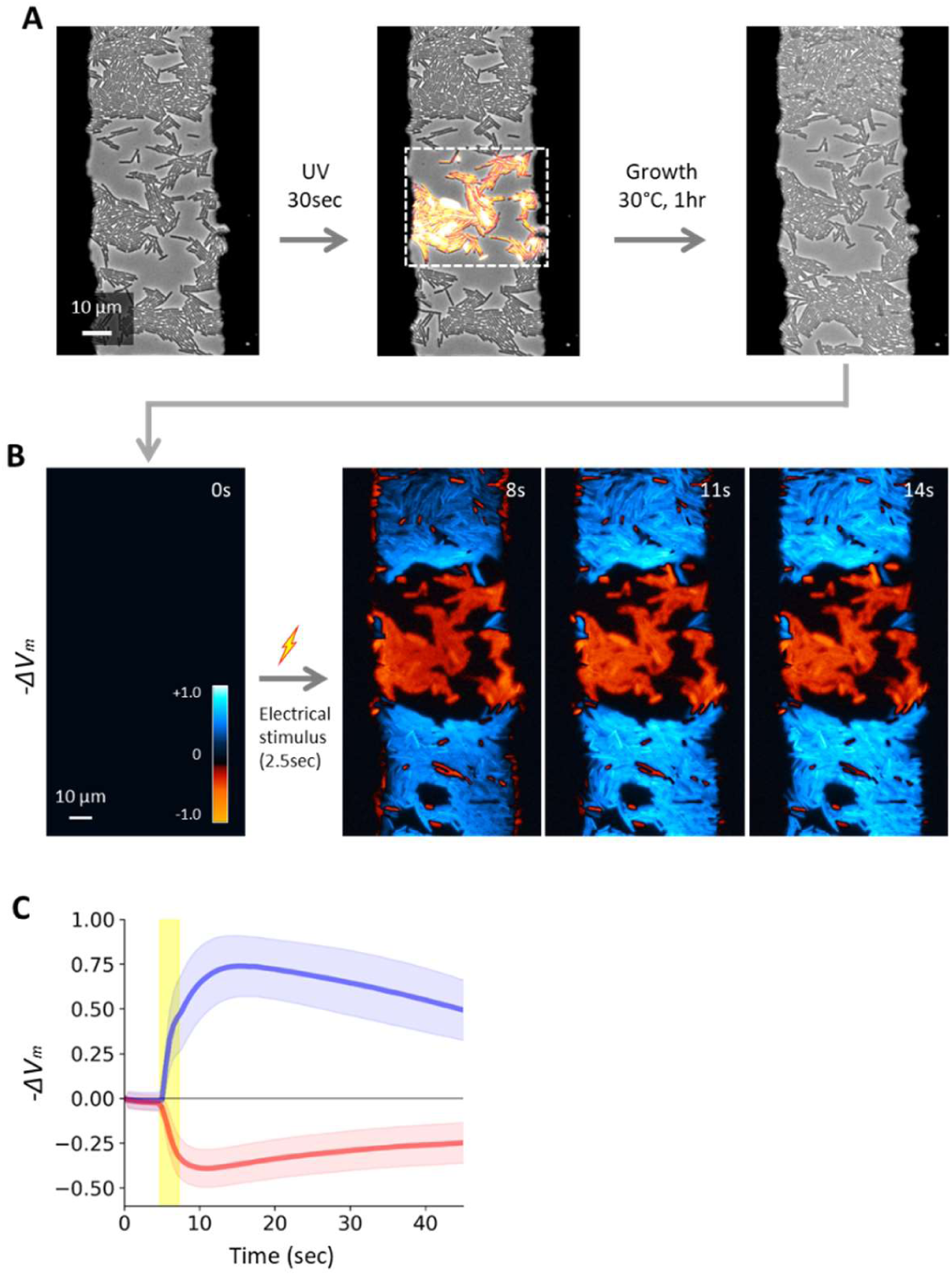
UV irradiation makes *B. subtilis* cells to response to an electrical stimulus in an opposite direction. A) Phase-contrast microscopy image shows wildtype *B. subtilis* cells within the electrode gap. A rectangular region indicated by the dashed line within the field-of-view was irradiated by UV light. Growth was suppressed in the UV irradiated region, while cells outside of UV-irradiation region replicated. B) The region shown in panel A was treated with an electrical stimulus. −ΔVm was calculated from ThT fluorescence (log(*F*_*ThT*_*/F*_*ThT,R*_)) and shown with the colormap in the panel. To an identical electrical stimulus, unperturbed cells hyperpolarized (blue) and UV-irradiated cells depolarized (red). C) Mean (thick lines) and Standard deviation (shaded color) of −ΔVm for cells in unperturbed (blue) and UV irradiate (red) regions.

### A mathematical model suggests that the response differentiation is due to the shift in resting membrane potential

To gain conceptual understanding of the observed distinct responses to an identical electrical stimulus, we used a mathematical framework based on FitzHugh-Nagumo (FHN) neuron model. The FHN neuron model, originally published over half a century ago (45), is one of the most paradigmatic models in neuroscience due to its mathematical simplicity and richness for capturing complex behaviors (46). We extended the FHN neuron model to bacterial electrophysiology while retaining its mathematical simplicity (see *SI text* for details). Briefly, in our FHN bacteria model, we considered two parameters representing the resting-state membrane potential and K^+^ transmembrane gradient. Numerical simulations of the model showed that an external electrical stimulus causes hyperpolarization in proliferative cells, while the same stimulus produces a relaxation response from depolarization in inhibited cells (Fig. 3A and S12). This is because the direction of K^+^ flux (influx or efflux) differ depending on the resting-state membrane potential and transmembrane concentration gradient of K^+^ (Fig. 3B). According to our simulations, opening of K_+_ channels in proliferative cells results in K_+_ efflux following the concentration gradient, thus causes hyperpolarization. However, the same opening of K^+^ channels only lead to the relaxation from depolarization due to weaker transmembrane K^+^ gradient. This mechanistic insight from the simulations predicts that the shift in resting-state membrane potential is sufficient to alter the response dynamics to an electrical stimulus. This means that different classes of growth-inhibition treatments should also make cells to respond by depolarization.

**Figure 3.**
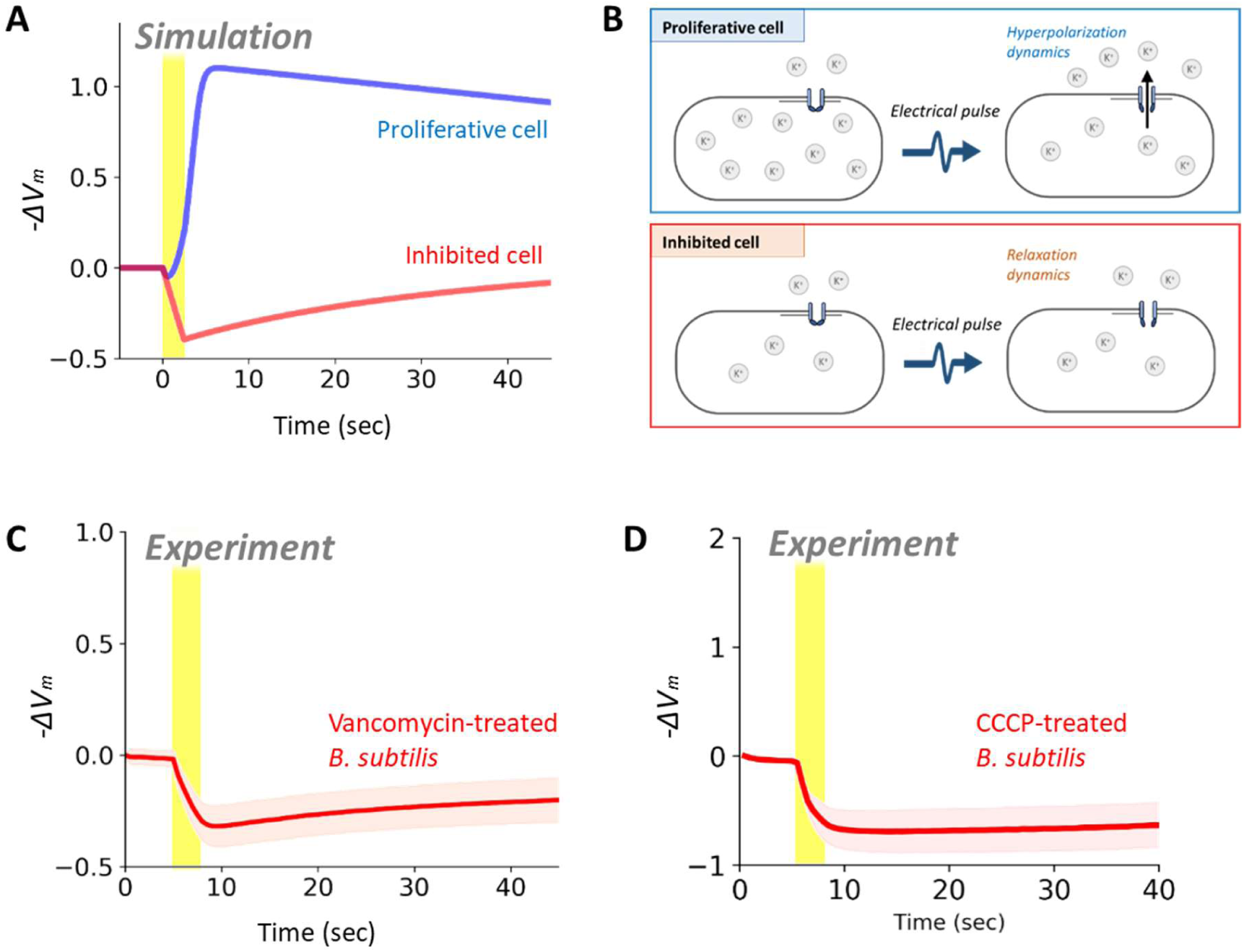
Shift in resting-state membrane potential is sufficient to describe the distinct responses between proliferative and inhibited cells. A) Numerical simulation result of the FHN bacteria model, corresponding to Fig. 2C. Despite being stimulated by an identical electrical stimulus, proliferative cells (blue) hyperpolarize and inhibited cells (red) depolarize. See SI text for model details. B) Illustration of biological mechanism of the response differentiation between proliferative and inhibited cells. C, D) Time trace of membrane potential change (−ΔVm) with *B. subtilis* cells exposed to (C) vancomycin or (D) CCCP. Shaded are standard deviation.

To examine this prediction from the model, we conducted the electrical stimulation experiment with the cells exposed to different classes of common growth-inhibition treatments; namely, an antibiotic vancomycin, a protonophore CCCP and a common antimicrobial agent ethanol. As predicted by the model, vancomycin-treated cells exhibited a clear depolarization in response to an exogenous electrical stimulus (Fig. 3C), as opposed to the hyperpolarization response seen in unperturbed cells (Fig. 1C). Experimental tests with the cells treated with ethanol or CCCP also showed depolarization response (Fig. 3D and S14). These results strongly support the conclusion drawn from the FHN model that proliferative and inhibited cells produce opposite membrane-potential dynamics, despite cells being treated by an identical electrical stimulus. Conversely, this finding suggests that the proliferative and inhibited cells can be distinguished based on their response to an exogeneous electrical stimulus.

### Selective antibiotics enables classification of coliforms in a mixed culture

The above observations and understanding raised the possibility of using the electrically induced membrane-potential dynamics for differentiation of species in a mixed culture based on the sensitivity to selective antibiotics. This process would find a wide range of applications where detection of proliferative bacteria is crucial. For proof of concept, we explored this possibility using a co-culture of *E. coli* (Gram-negative) and *B. subtilis* (Gram-positive). Our hypothesis was that the above approach of measuring electrically induced membrane-potential dynamics allows for the rapid differentiation of *E. coli* and *B. subtilis* when combined with exposure to vancomycin. This is based on the fact that vancomycin inhibits the cell-wall synthesis of Gram-positives but is largely inactive to Gram-negatives due to their outer membrane barrier (47).

To test this hypothesis, we co-cultured fluorescently labeled strains of *E. coli* and *B. subtilis* expressing mCherry and YFP under IPTG inducible promoters for *E. coli* and *B. subtilis*, respectively. Multi-channel fluorescence imaging exhibited distinct signals for *E. coli* and *B. subtilis* (Fig. 4A). The fluorescence of ThT showed lower intensity with *E. coli* compared to *B. subtilis* (Fig. 4B). Although the exact reason for this is unclear it may suggest that there is a difference in stability or fluorescence yield of ThT between these two species. This difference in intensity level highlights an advantage of our approach focusing on the response dynamics rather than being reliant on the initial affinity of the cells for the dye. After an hour-long exposure to vancomycin, the co-culture was treated with an external electrical stimulus. The result revealed distinct patterns of membrane potential dynamics for *E. coli* and *B. subtilis* cells treated with vancomycin. Specifically, the electrical stimulation causes *E. coli* to become hyperpolarized, while depolarizing *B. subtilis* (Fig. 4 C, D and Fig. S16). Similar patterns were observed with mono-cultures of *E. coli* or *B. subtilis* (Fig. S15). We also confirmed that vancomycin is active on *B. subtilis*, but not on *E. coli* (Fig. S13). The histogram of individual-cells response revealed distinct distributions between *E. coli* and *B. subtilis* (Fig. 4E). These results thus demonstrate that the approach focusing on electrically induced membrane-potential dynamics allows for rapid differentiation of *E. coli* and *B. subtilis* in a mixed culture.

**Figure 4.**
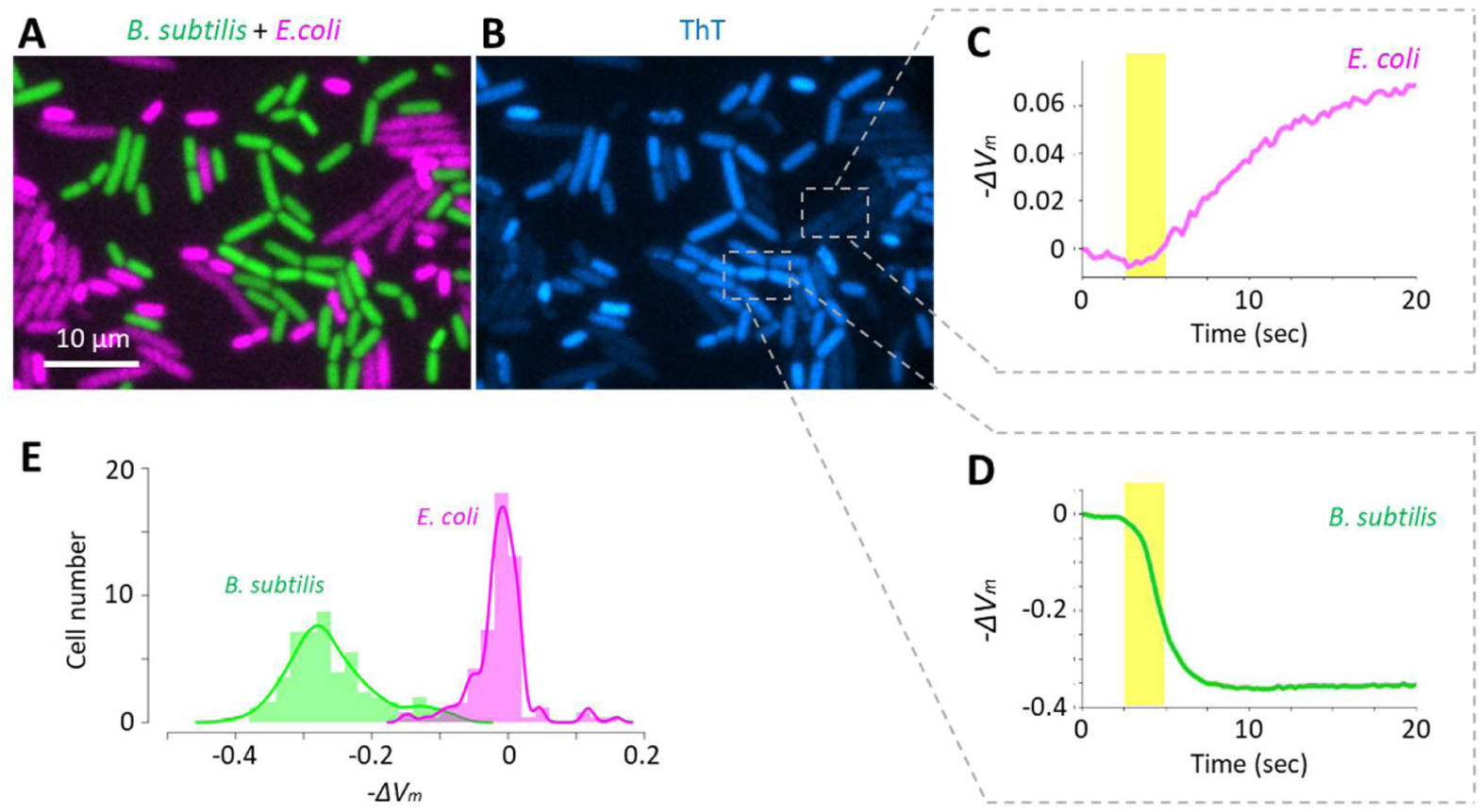
*B. subtilis* and *E. coli* cells can be differentiated based on their response to an electrical stimulus. A) Microscopy image showing the co-culture of fluorescently labeled *E. coli* (magenta) and *B. subtilis* (green) cells treated with vancomycin. B) ThT fluorescence image at the corresponding region. C, D) Time trace of −ΔVm, calculated from ThT fluorescence intensity, of the regions defined in panel B. E) Histogram of −ΔVm at 10 sec after electrical stimulation of *B. subtilis* and *E. coli* shows clear differentiation of peaks with the presence of vancomycin.

## Discussion

Our hope is that the experimental setup we have developed and demonstrated in this work will encourage more microbiologists to consider bacterial electrophysiology in conjunction with their physiological processes of interest. In addition to mechanosensation and biofilm dynamics, membrane potential is closely associated with important microbiological processes including persister formation and antibiotics resistance (48–51). This is an important societal challenge since antimicrobial resistance is rising. Our device will provide a new toolkit to identify the molecular-mechanism involved in this relation. It would also be useful to systematically analyze whether externally altering membrane potential using electrical stimulation impacts the formation of persisters. Furthermore, our device could be used for single-cell investigation of metabolic electron flow, which would be particularly important for metabolic engineering (52–54). Beyond these phenomena known to be related to membrane potential, we speculate that bacterial electrical signaling may play roles in many more physiological processes than previously realized. The uses of exogeneous electrical stimuli should unlock opportunities to gain new insights regarding bacterial electrical signaling mediated by membrane potential dynamics. In parallel to fundamental research, we envision that the use of applied electrical stimuli in the context of bacterial electrophysiology will open the door to future developments into a novel approach for real-time electrical control of bacterial functions.

Our finding, that proliferative capacity alters cellular response to electrical stimuli, provides insights into bacterial electrical signaling. For example, it was shown that metabolic stress, namely glutamate limitation, triggers biofilm electrical signaling (3). Our findings now suggest that metabolic stress is not only capable of initiating electrical signaling but also alters the cellular response to this signaling. This additional layer of interaction between metabolic activity and electrical signaling can form a feedback loop that allows emergence of complex behavior. Characterizing this feedback interaction and examining the impacts of electrical stimulus to biofilms are important future research tasks. In *E. coli* mechano-sensation, electro-physiological response to mechanical stress was shown to be heterogeneous, although the source of this heterogeneity remains unclear (1). If the response to electrical stimuli is intrinsically linked to proliferative capacity, then our discovery may suggest that the endogenous heterogeneity in proliferative capacity can be a source of this heterogeneity. It will be interesting to examine this possibility using single-cell microscopy analysis combined with an applied exogeneous electrical stimulus.

Our simulation and experiments suggested that the distinct response dynamics between proliferative and inhibited cells are due to a shift in resting membrane potential. Since the maintenance of membrane potential accounts for a major fraction of cellular energy consumption (30), it seems plausible that different types of metabolic and environmental stress would all ultimately lead to imbalance in transmembrane ion gradient at resting state. For instance, the Ktr potassium uptake system in *B. subtilis* and the Kdp potassium uptake system in *E. coli* are both ATP-driven (55,56). This means that maintaining the resting-state membrane potential requires constant consumption of ATP in order to keep the intracellular K^+^ level up to two orders of magnitudes higher than the outside. When an electrical stimulus opens voltage-gated K^+^ channels, the flux of K^+^ through the channel follows the electrochemical gradient of K^+^, which indicates that hyperpolarization due to K^+^ efflux occurs only when intracellular K^+^ concentration is significantly greater than the extracellular level. This effect was accounted by the parameter *k*_*K*_ in our FHN bacteria model (see *SI text*). In the future studies, we will quantitatively analyze the dynamics of other electrophysiologically important ions, such as Ca^2+^ and Cl^-^, and determine their contributions to the membrane potential dynamics in bacteria. Biological and biophysical characterization of the dynamics of transmembrane gradients of different ions and their corresponding channels will form the fundamental basis to our understanding of bacterial electrical signaling. Another important area of research is to identify the molecular mechanism by which electrical stimuli open the YugO potassium channel.

Finally, our findings offer a novel approach for rapid detection of proliferative bacteria without the need of observing actual proliferation. Most importantly, the focus on response dynamics minimizes the requirement of calibrations for differentiation of proliferative and inhibited cells. This contrasts with the currently available techniques relying on single-time point measurements which could be affected by many physiological states in various ways such as species, growth phases and media (33). It is also worth noting that our approach could detect proliferative cells within a minute after an electrical stimulation, as opposed to the typical duration of 12-48 hours required by conventional culture-based detection methods (57). This is an attractive feature since it could accelerate the examination of antimicrobial agents and diagnosis in medical sector and enable efficient quality-control in the water, pharmaceutical, food and beverage industries. The growing demand for fast identification of live bacterial cells has been driving the developments of novel technologies for rapid bacterial detection (57). Our findings provide a novel strategy with a unique capability to differentiate UV-damaged cells from healthy cells. In further studies, we will examine if this approach enables the detection of medically and industrially relevant bacterial species for timely diagnosis. If applied widely, this approach of using membrane-potential dynamics and exogenous electrical stimuli could bring great societal benefits by accelerating the detection of proliferative bacteria and determination of their sensitivity to antimicrobial agents.

## Methods

### Strains and Growth conditions

*Escherichia coli* and *Bacillus subtilis* cells were routinely grown in Lysogeny broth (LB) or on LB agar (1.5 % (w/v)) plate. The reporter and mutant strains used in this study are listed in Table S1. For electrical stimulation experiments, a colony from LB agar plate was inoculated into liquid LB and subsequently incubated at 30°C with aeration (200 r.p.m.) to OD600 ≈ 1.5. Cells were then resuspended in MSgg (minimal salts glutamate glycerol) media (58): 5 mM Potassium Phosphate (pH 7.0), 10 mM MOPS (pH 7.0), 2 mM MgCl2, 700 μ M CaCl2, 50 μ M MnCl2, 100 μ M FeCl3, 1 μ M ZnCl2, 2 μ M thiamine-HCl, 0.5 % (v/v) glycerol, 0.5 % (w/v) monosodium glutamate. Note that the MOPS concentration is reduced by 10-fold from the original receipt of MSgg in order to suppress electrolysis of the media. After an hour incubation in liquid MSgg, cells were inoculated onto MSgg LMP (low-melting point) agarose pads containing 10 μ M Thioflavin T (ThT) (Sigma-Aldrich). With experiments focusing on mixed culture, 5 μ g/ml vancomycin hydrochloride, 1 mM NH4Cl and 0.25% (w/v) glucose were supplemented to the MSgg LMP-agarose pads. Pads were prepared as described previously (59). Briefly, LMP agarose (Formedium, Bacteriological Granulated Agar) was dissolved in MSgg and left to solidify between two 22 mm x 22 mm cover glasses (Fisher Scientific) for 10 minutes at room temperature. When stated, final concentration of 1 % (v/v) ethanol or 100 μ M CCCP was supplemented to MSgg liquid and MSgg agarose pads. For the measurements of intracellular K^+^, APG-2 (Abcam PLC), instead of ThT, was supplemented to MSgg agarose pad at the final concentration of 2 μ M. The solidified agar was cut into approximately 5 mm x 5 mm pads. 2 μ l of bacterial liquid LB culture (OD600 ≈ 1.5) was inoculated onto each pad. Pads were then placed on the gold-coated glass bottom dish for microscopy.

For the construction of *E. coli* K12 pGEX6P1-mCherry strain, pGEX-6P1 plasmid (GE Healthcare) was digested with EcoRI and NotI restriction enzymes (New England Biolabs). The mCherry gene was amplified by PCR using PrimeSTAR Max DNA Polymerase (Takara Bio) using the primers AP609 (5’-CCCCTGGGATCCCCGGAATTCATGGTGAGCAAGGGCGAG-3’) and AP610 (5’-AGTCACGATGCGGCCGCTCGAGTTTAGCACTTGTACAGTTCGTCCATG-3’). The PCR product was assembled together with the digested pGEX-6p1 by the Gibson Assembly using Gibson Assembly Master Mix kit (New England Biolabs), and was transformed into competent *E. coli* K12 cells. The competent cells were prepared using Mix&Go Competent Cells kit (Zymo Research). The sequence of the assembled plasmids was confirmed by Sanger sequencing (Source BioScience) and aligned using Benchling (https://benchling.com/).

### Relay circuit

Components shown in Fig. S3 were mounted through-hole onto a solderable breadboard with copper tracks printed onto one side. Connections between components were accomplished by soldering jumper wires of appropriate length to the appropriate position on the copper track. A track breaking tool was used to isolate the components which should not be connected but were on the same track. The components were assembled as shown in the circuit diagram (Fig. S3). The trigger input from scientific cMOS camera (Orca Flash 4.0, Hamamatsu) was connected to the circuit through a BNC (Bayonet Neill-Concelman) connector soldered onto the circuit board. The signal pin from the BNC connector was further connected to an analogue pin on an Arduino UNO R3 (Arduino, <u>arduino.cc</u>) to record when the voltage output from the camera attached to the microscope. A counting loop was implemented in Arduino script to count camera exposures to coordinate the electrical stimulate and sequential imaging. At the programmed time in the imaging sequence a digital pin is set ‘HIGH’ to provide power to an electro-mechanical relay. This switches the relay throw from ground to the input signal from the arbitrary function generator (Tekronix). All counter electrodes are permanently grounded through a common ground connection.

### Constructing Bespoke Electrode Dishes

Stainless steel disks of 50 μ m in thickness and 39 mm in diameter were manufactured by Laser Micromachining Ltd. with a negative electrode design and used as coating masks. A negative mask was placed into a glass-bottom dish (HBST-5040, Willco Wells). The glass-bottom dish with a mask in place was then attached to 40 mm stainless steel washers on a silicon wafer using several 5 mm neodymium magnets. Once attached, the silicon wafer was mounted in an Electron Beam Physical Vapor Deposition System (A custom built model II2000EB, Scientific Vacuum Systems); a 20 nm titanium adhesion layer and a 100 nm gold conductive layer were deposed onto the dish, with the mask permitting the electrode design to be applied. Copper wire was soldered to corresponding connections on a 10 pin IDC (Insulation-displacement connector, RS Components) at channels corresponding to the electrodes. Small holes were made through the plastic rim of the dish, with the copper wire being threaded through until all were in their corresponding positions. The IDC was then fixed in position using two-part waterproof adhesive (Araldite Rapid, RS Components). Once the IDC was fixed in place the wires were attached to their corresponding electrode terminals with aqueous graphite conductive adhesive (Alfa Aesar), then left overnight at room temperature to set to reduce resistance of the adhesive. Following this, the graphite connections were covered with Araldite adhesive to protect them from moisture degradation.

### Electrical Stimulation

Application of electrical stimulation was accompanied with time-lapse imaging with 2 fps (frames per second) for 1 minute. An alternating current (AC) signal (0.1 kHz; 3 V peak-to-peak (−1.5∼+1.5 V)) was generated using an Arbitrary Function Generator (Tektronics) and connected to a series of relays, each corresponding to an electrode on the gold-coated dish (Fig. S2). The camera trigger was connected to Arduino UNO R3 in the relay circuit to control the timing of electrical stimulation; upon counting 10 camera exposures, the relay to the electrode being imaged would open for 2.5 sec, applying electrical stimulation to the electrode while simultaneously imaging.

### Time-lapse Microscopy

The membrane potential dynamics and growth of individual cells were recorded using an inverted epi-fluorescence microscope, DMi8 (Leica Microsystems), operated by MetaMorph (Molecular Devices). The microscope is equipped with an incubation chamber (Pecon, i8 Incubator) which maintained the temperature at 30°C throughout the experiments. Prior to microscopy experiments, the chamber was set to 30°C for at least 3 hours, and samples were placed in the chamber for an hour. For all observations, a 100x objective lens (NA=1.3, HCX PL FLUOTAR, Leica) was used and images were taken with scientific cMOS camera ORCA-Flash 4.0 v2 (Hamamatsu Photonics). Cell growth was monitored using phase contrast (exposure time: 100 msec). ThT fluorescence was detected using a single-band filter set consisting of Ex 438/24 nm, Em 483/32 nm, and Dichroic mirror 458 nm (Semrock), with exposure time of 150 msec. For the mixed culture experiment, YFP was detected using a filter set consisting of Ex 509/22, Em 544/24, and Dichroic mirror 526 (Semrock). mCherry was detected using a filter set consisting of Ex 554/23, Em 609/54, and Dichroic mirror 573 (Semrock). The exposure time for the imaging of both YFP and mCherry was 300 msec.

For irradiation to UV violet light, cells were irradiated by UV-Violet light for 30 sec using the inverted microscope DMi8 (Leica Microsystems) and the LED light source, SOLA SM II Light Engine (Lumencor), with an excitation filter 400/16. Field Diaphragm of the microscope was used to irradiate only a specific region of the field of view. Before electrical stimulation, a one-hour growth period was allowed during which cells were observed using phase contrast microscopy to ascertain the effects of UV-V light. For ethanol experiments, MSgg containing 1% (v/v) ethanol was used instead of Msgg.

### Image analysis

Region of interest (ROI) was registered for individual cells and the fluorescence intensity over time for each ROI was measured using Fiji/imageJ. The results were saved in csv (comma-separated values) and imported to python. Based on the Nernst equation (*See SI text* for details), −ΔVm was calculated by taking the logarithms of the ratio between intracellular ThT fluorescence intensity (*Fi*) and the intensity at the resting state (*Fi,*_*R*_). For measurement of cell elongation rate, ROI was registered for individual cells and the cell size over time for each ROI was measured. The results were saved in csv and imported to MATLAB. The elongation rates were determined as the slope of fitted straight lines. Numerical calculations were performed in python using scipy (scipy.org) and numpy (numpy.org). Graphs and plots were generated using the python data-visualization packages matplotlib (matplotlib.org) and seaborn (seaborn.pydata.org). Scripts were written using Jupyter Notebook (jupyter.org/).

## Acknowledgements

We thank R Jeanneret for his help with initial prototyping; M Crouch for his assistance with Vapor Deposition System; O Back for his help with the simulation; P Schäfer, DY Lee, GM Süel, M Kittisopikul, A Mica and the members of the Asally lab for their comments to the manuscript; A Prindle, GM Süel and A Pires-daSilva for generously providing bacterial strains. This study was supported by funding from University of Warwick, Royal Society Research (RG150784) and Innovate UK (15944) to MA, BBSRC/EPSRC grant to Warwick Integrative Synthetic Biology Centre (BB/M017982/1) and EPSRC/BBSRC Synthetic Biology Centre for Doctoral Training to MD (EP/L016494/1).

## Author Contributions

JPS and MA conceived the project and developed the device. JPS, CLAE, MJG, DM and MD prepared materials for experiments and tested the device. CLAE, JPS, MJG and DM conducted experiments. CLAE, MJG, DM and MA analyzed the results. YH and MA developed the model and performed simulations. MA wrote the manuscript with the inputs from all authors.

## Supplementary Figures

**Figure S1.**
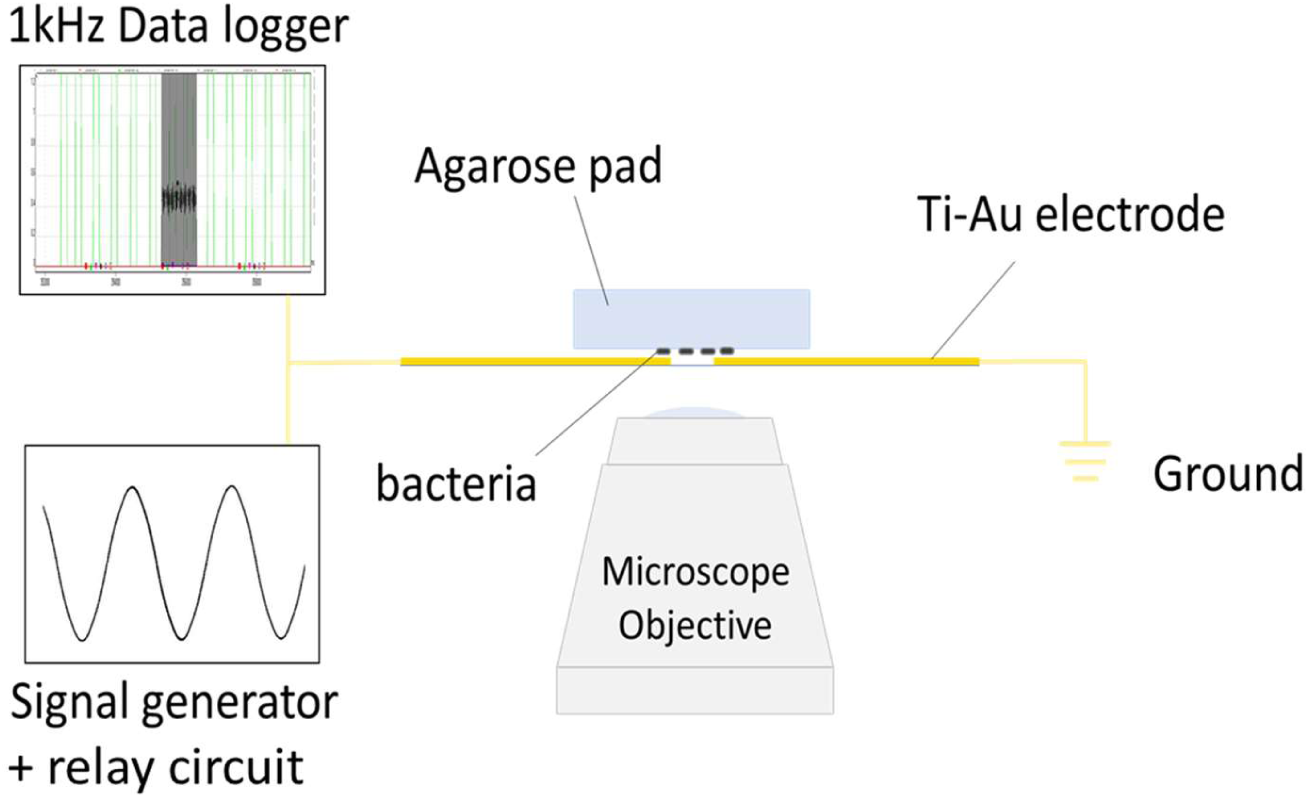
Schematic diagram of the experimental setup. The Ti-Au electrode coated dish (Fig. 1A) is connected to a data logger, signal generator and a relay circuit. Bacteria are inoculated on an agarose pad and placed on electrode-coated dish.

**Figure S2.**
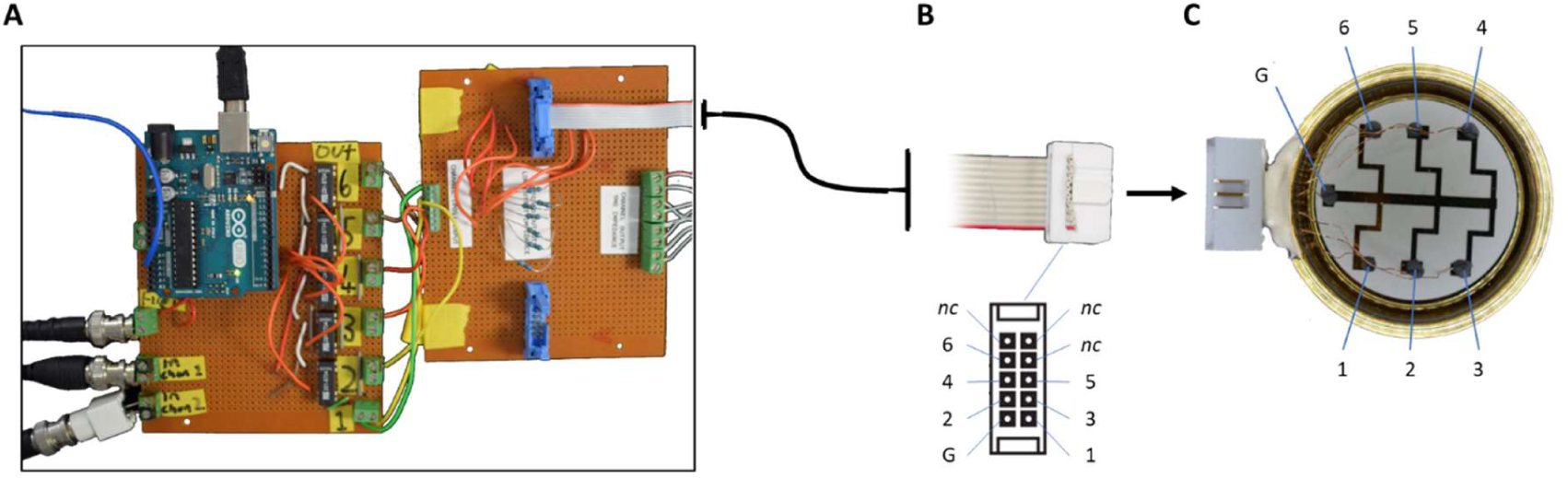
Schematic images of relay circuit and electrode-coated dish. A) Photograph of the custom-built relay circuit with Arduino UNO R3. B) The pin layout of 10-pin IDC output was used to connect the relay circuit and dish. Numbers correspond to each position shown in panel C. nc is not-connected. C) The electrode-coated dish image with the labels of corresponding terminals of each pin. This provides an individual control of firing electrodes.

**Figure S3.**
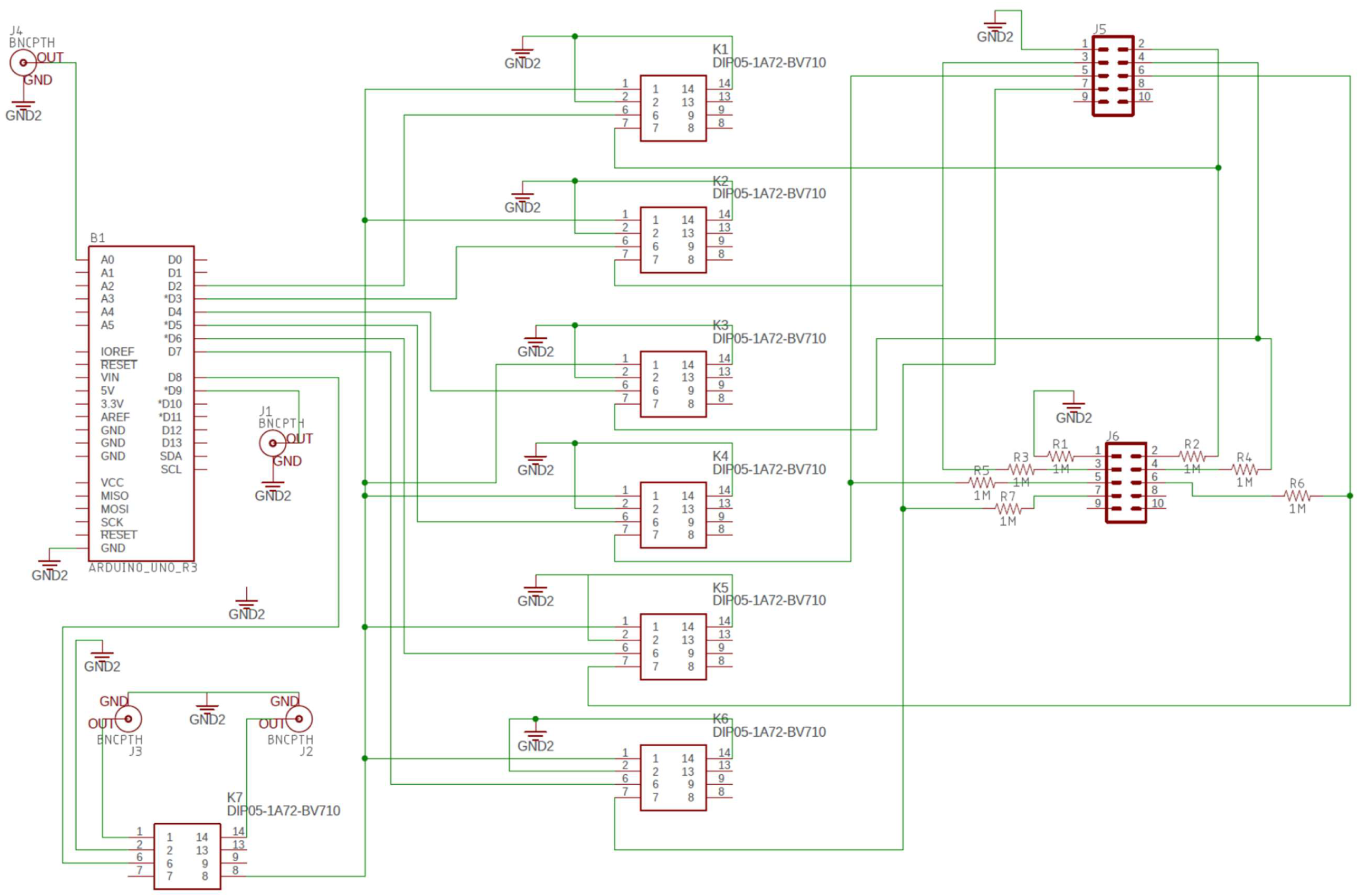
Schematic diagram of relay circuit. Schematic diagram of the relay circuit with Arduino UNO. K1-6 correspond to channels shown in Fig. S2.

**Figure S4.**
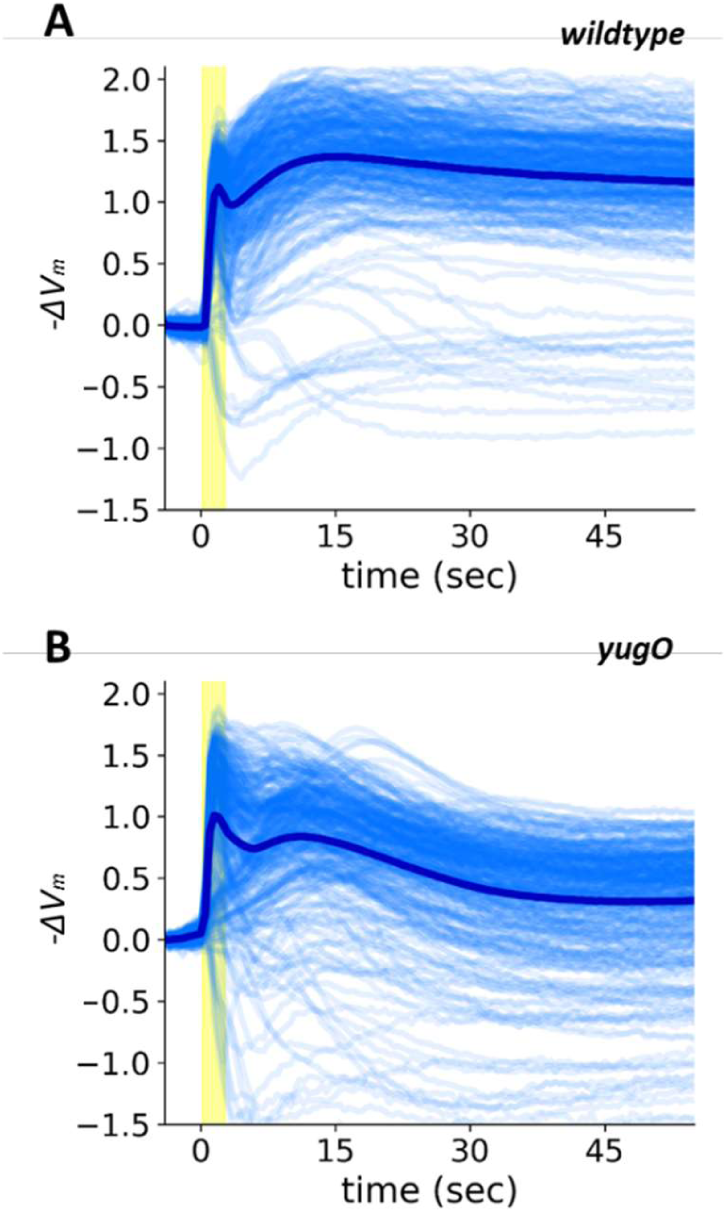
Single-cell time traces of membrane-potential response to an electrical stimulus. A) wildtype and B) *yugO* strain. −ΔVm was calculated from ThT and plotted for individual cells.

**Figure S5.**
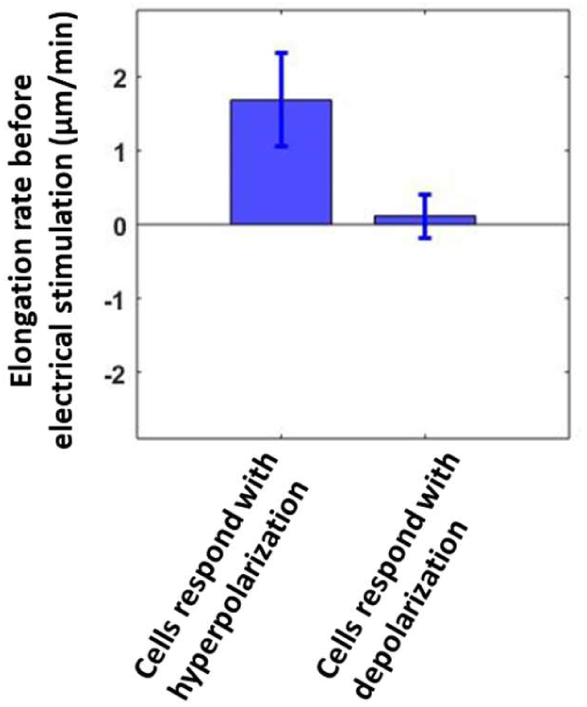
Growth of the cells respond with depolarization is halted before electrical stimulation. Mean elongation rates before electrical stimulation were quantified for the individual cells. (n = 15 cells for each). Error bars are standard deviation.

**Figure S6.**
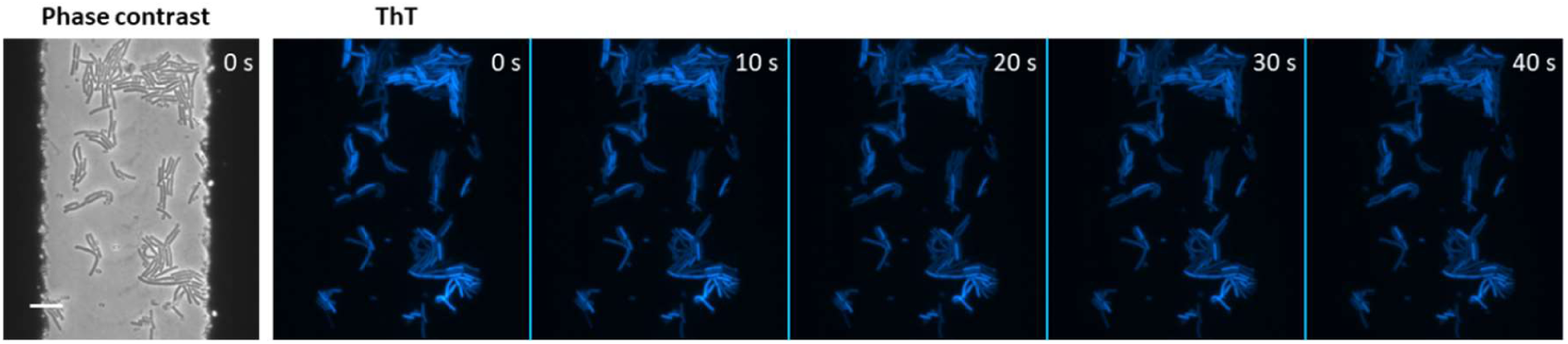
Resting-state membrane potential is static. Film-strip images of ThT fluorescence without electrical stimulation. No significant changes were observed without electrical stimulation.

**Figure S7.**
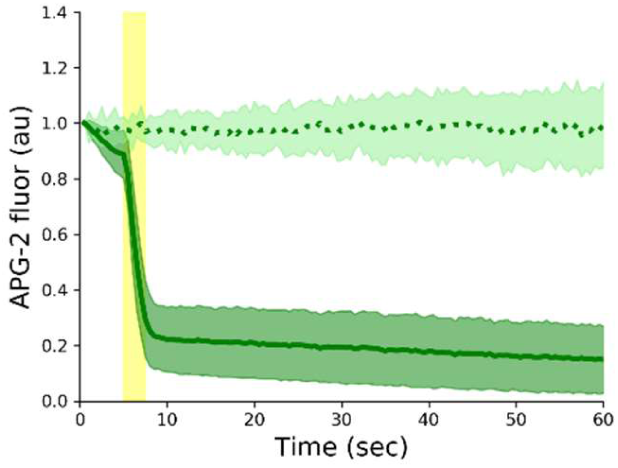
Electrical stimulation causes K+ efflux. Mean intracellular potassium levels measured with APG-2 AM with and without electrical stimulation. Solid green line is the mean APG-2 intensity with electrical stimulation (indicated in yellow). Dashed light green line is the mean APG-2 intensity without electrical stimulation. Shaded areas are standard deviation. (n = 60 cells for each).

**Figure S8.**
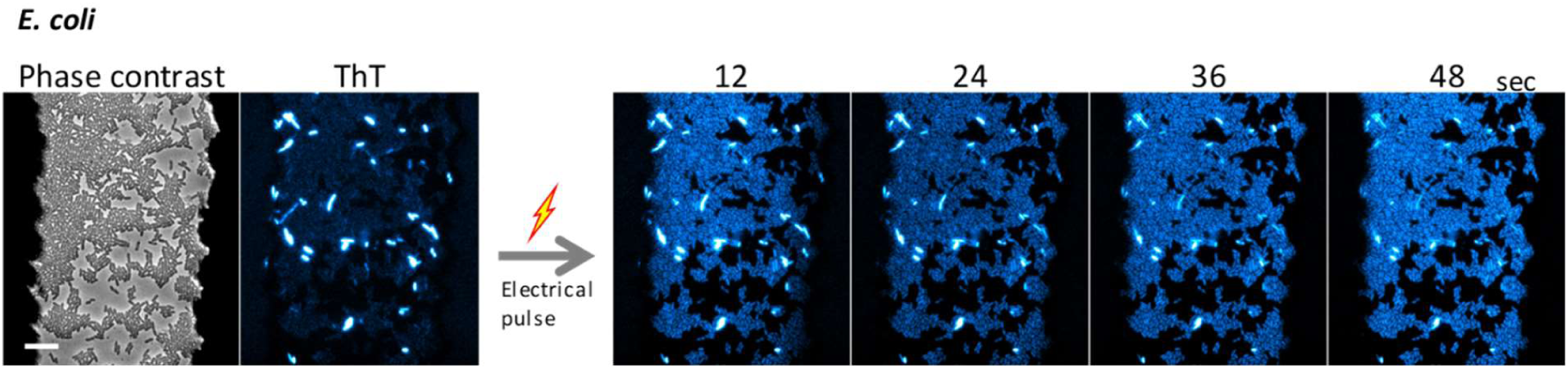
*E. coli* cells also exhibit hyperpolarization response to an electrical stimulation. Film-strip images of ThT fluorescence in *E. coli* before and after an electrical stimulation. Majority of cells hyperpolarize, while a small subpopulation of bright cells at resting state depolarize in response to an electrical stimulation.

**Figure S9.**
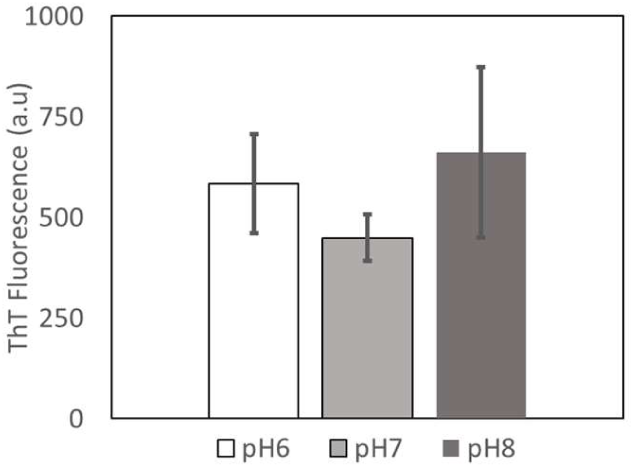
Mean ThT fluorescence intensity of *B. subtilis* cells at different pH are similar. ThT intensity was measured at single-cell level at different pH (n = 60 cells for each). Error bars are standard deviation.

**Figure S10.**
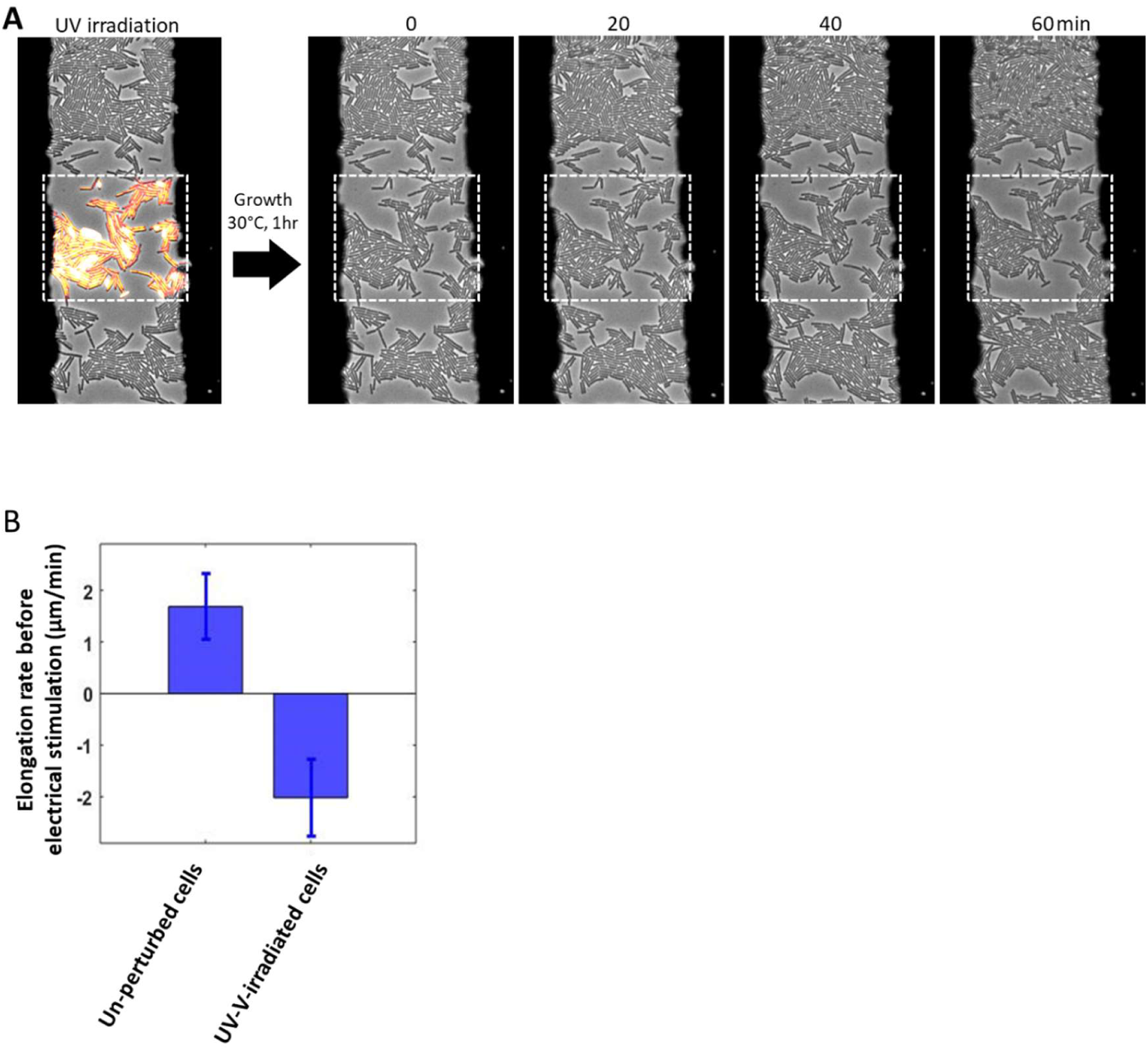
UV-V irradiation inhibit the growth of cells. A) Growth suppression by UV-V irradiation was confirmed by phase-contrast time-lapse microscopy. The film-strip images correspond to Fig. 2A. B) Quantification of elongation rates for unperturbed and UV-V irradiated cells (n = 15 cell for each).

**Figure S11.**
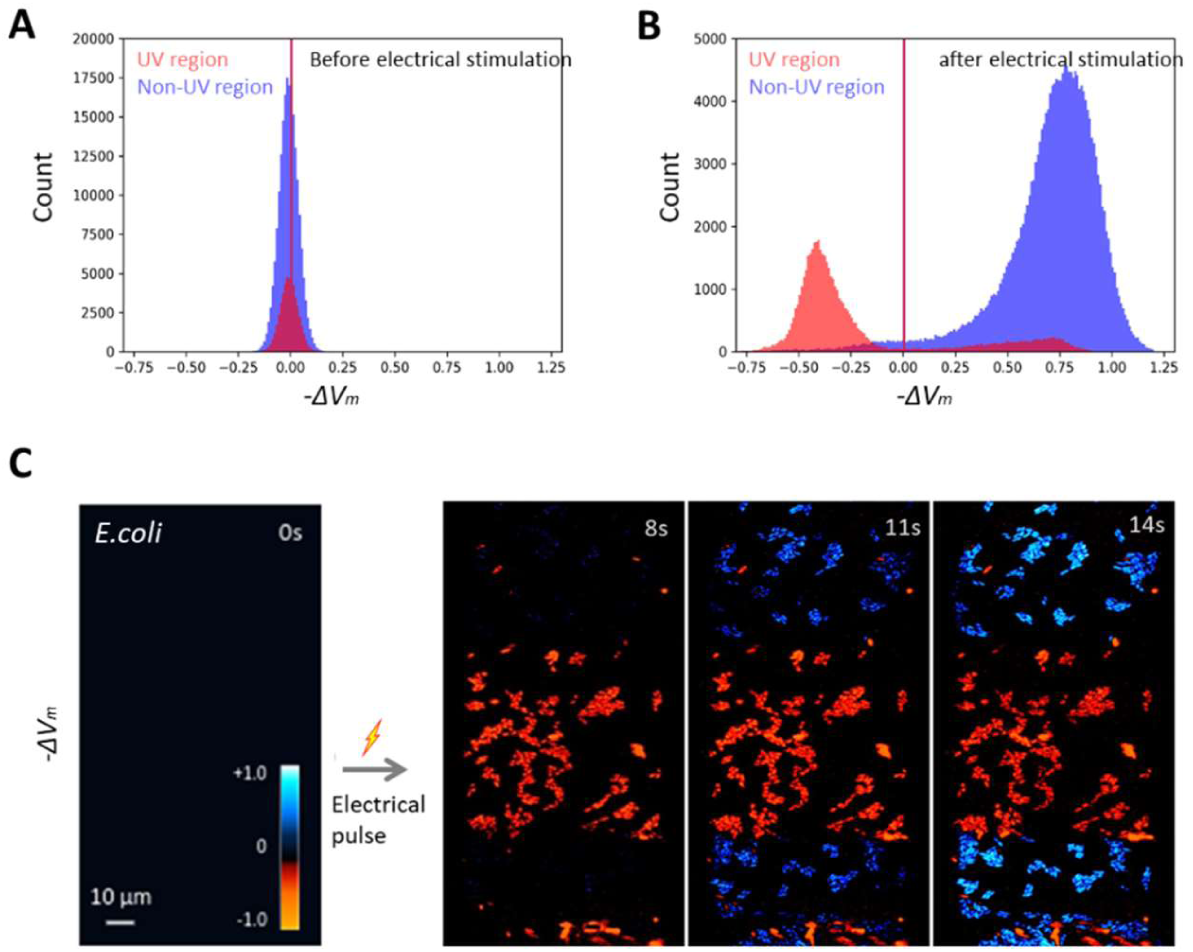
Response differentiation between unperturbed and UV-V irradiated cells. A, B) Histogram of the membrane potential A) before and B) after electrical stimulation. Blue and red are unperturbed and UV-irradiated regions, respectively. C) Membrane potential response dynamics of *E. coli* cells unperturbed and UV-irradiated cells.

**Figure S12.**
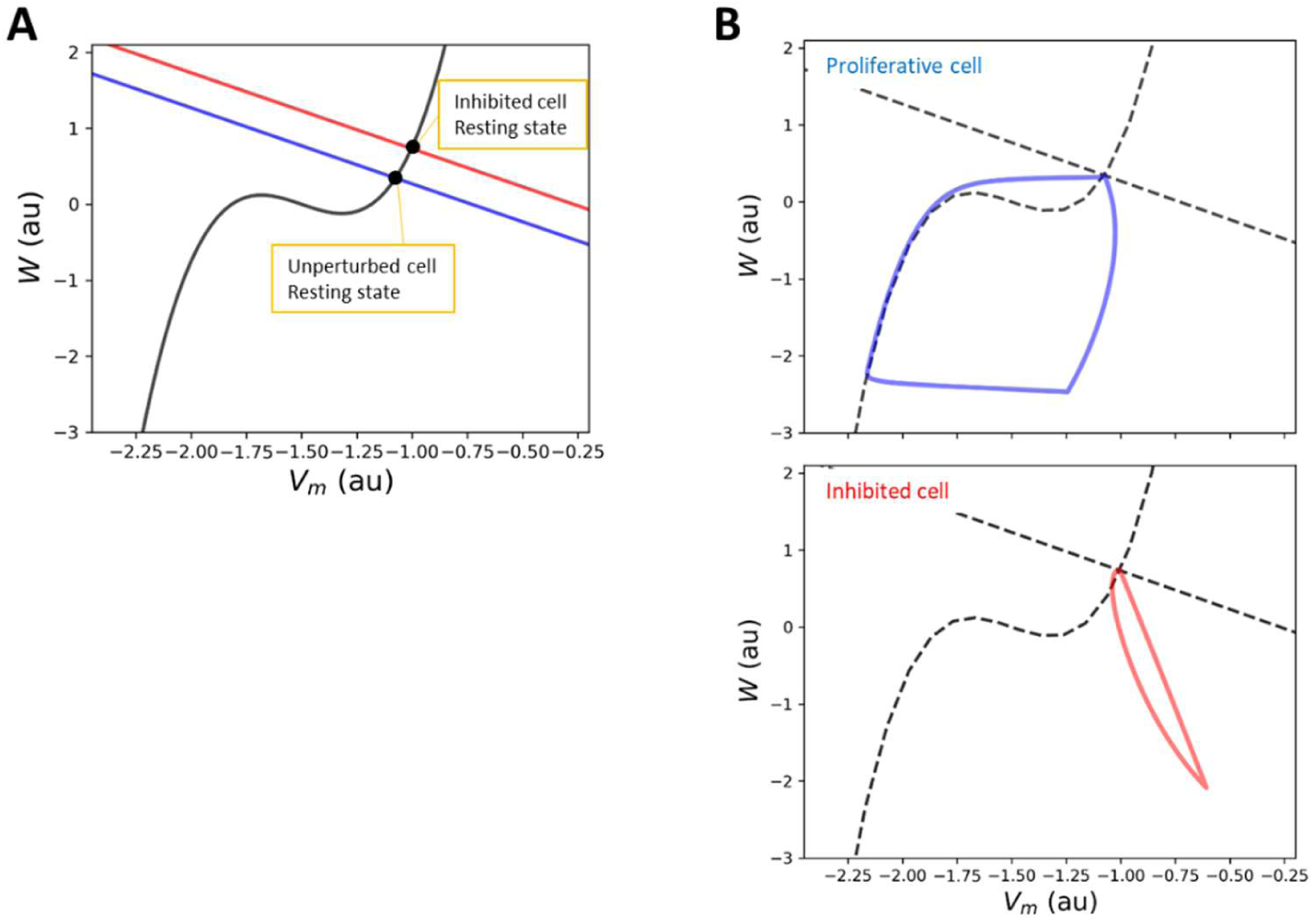
Simulations of the FHN bacteria model in Vm-W phase space. A) Nullcline of Vm is drawn in black. Nullclines of W with the parameters for proliferative and inhibited are drawn in blue and red, respectively. The stable equilibrium points of the system (resting state) are shown by black circles. B) Phase-space trajectories of the simulations for proliferative (blue) and inhibited (red) cells.

**Figure S13.**
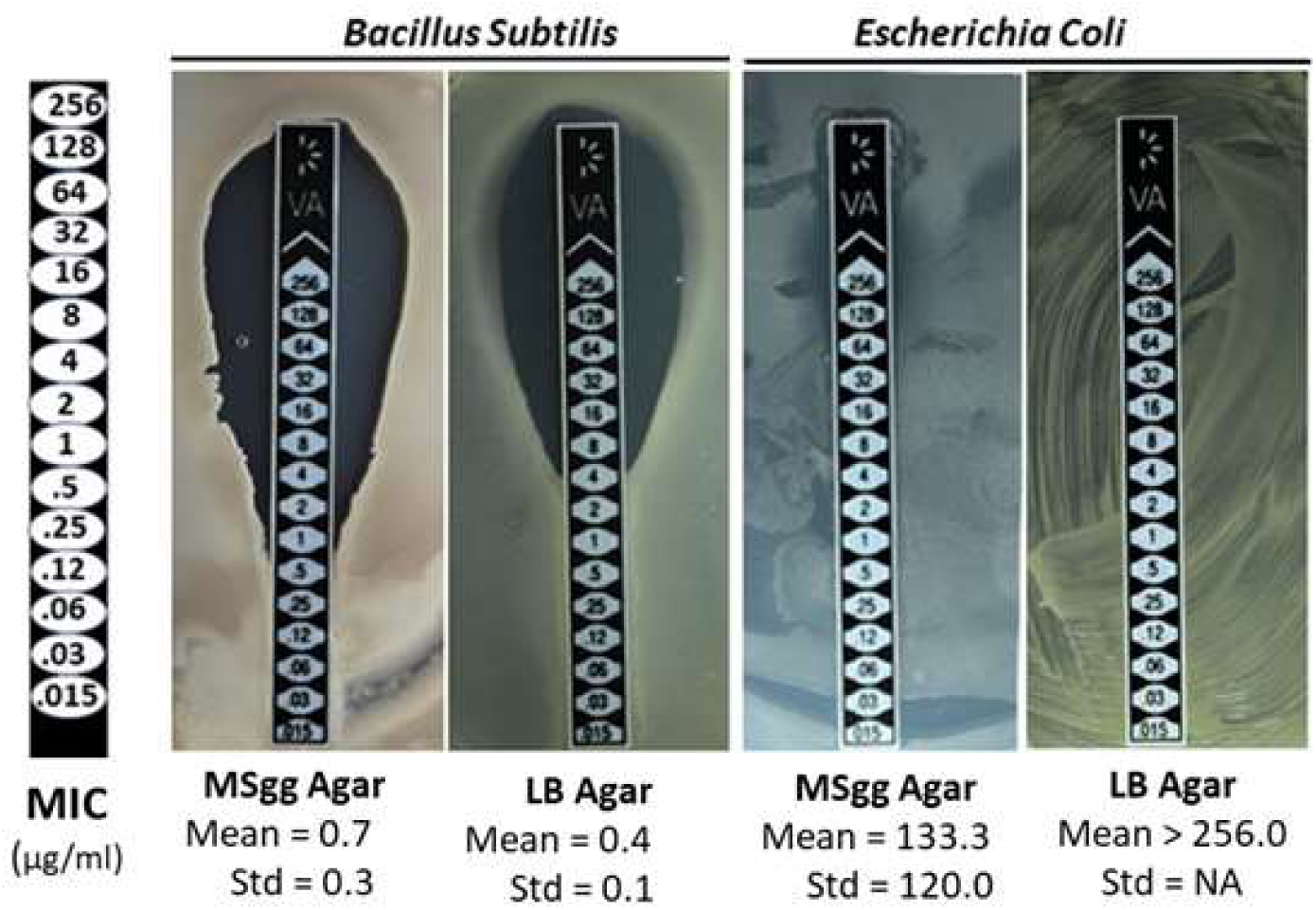
Minimal inhibitory concentration (MIC) for vancomycin against *B. subtilis* and *E. coli* on LB or MSgg agar plates. MIC was determined by reading the edge of clearance zone. The experiments were repeated three times.

**Figure S14.**
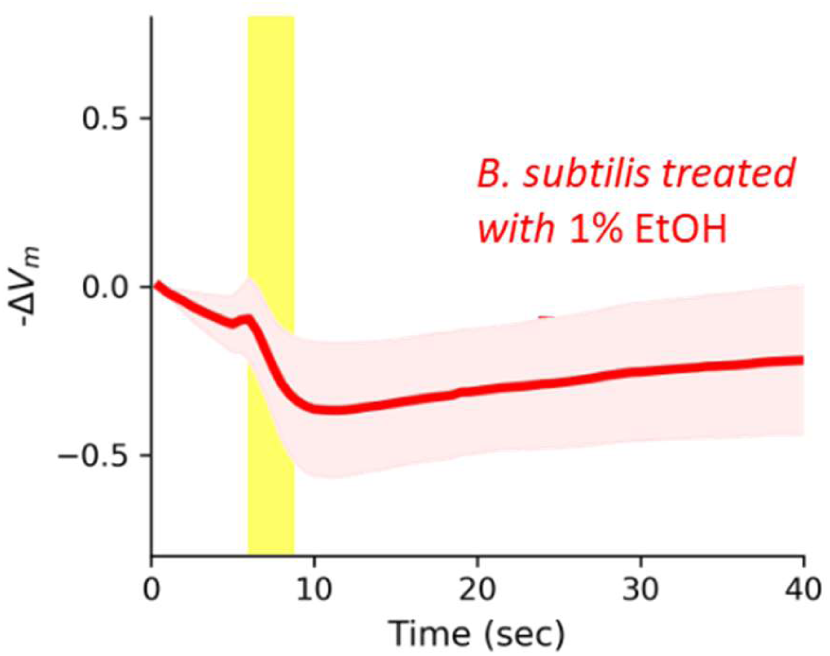
Cells treated with 1% Ethanol depolarize when stimulated by an electrical pulse. Membrane potential dynamics with an application of electrical stimulus (indicated in yellow). Solid line show the mean intensity and shaded is standard deviation. (n = 60 cells)

**Figure S15.**
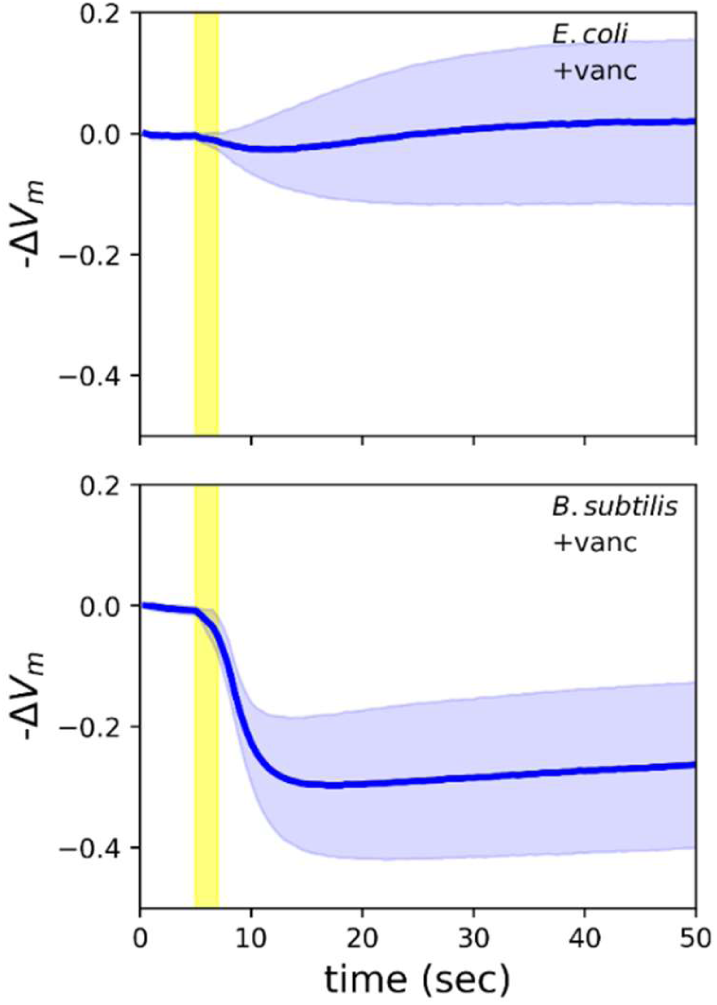
Membrane potential dynamics of *E. coli* and *B. subtilis* cells treated with vancomycin as mono-culture. Solid lines are mean intensity and shaded are standard deviation. (n = 20 cells for each).

**Figure S16.**
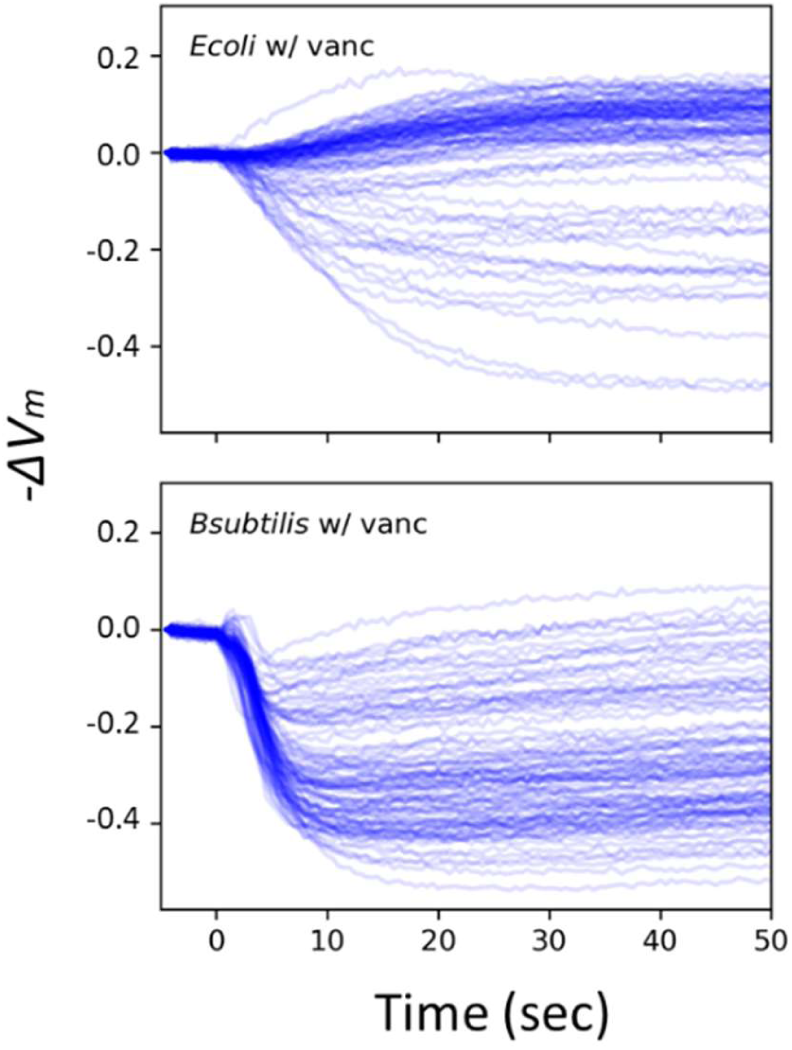
*E. coli* and *B. subtilis* can be differentiated based on their response dynamics to an electrical stimulus. Individual cells’ response dynamics to an electrical stimulation. The data correspond to Fig. 4E.

**Table S1.**

| Strain ID | Specie | Genotype/Plasmid | Source |
| --- | --- | --- | --- |
| MA-002 | *B. subtilis* | NCIB3610, wildtype | Lab Stock (57) |
| MA-020 | *B. subtilis* | NCIB3610, amyE:: *P<sub>hyperspank</sub>*-YFP, spec<sup>R</sup> | Gift from GM Suel (UC San Diego) (59) |
| MA-627 | *B. subtilis* | NCIB3610, yugO:: neo<sup>R</sup> | Gift from A Prindle and GM Suel (UC San Diego) (3) |
| MA-727 | *E. coli* | CGSC4401 EMG2 K12, wildtype | Gift from A Pires-daSilva (University of Warwick) |
| MA-946 | *E. coli* | CGSC4401 EMG2 K12, pGEX-6P1-mCherry | This work. |

## References

1. Bruni GN, Weekley RA, Dodd BJT, Kralj JM (2017) Voltage-gated calcium flux mediates Escherichia coli mechanosensation. Proc Natl Acad Sci 114(35):9445–9450.

2. Lee DD, Prindle A, Liu J, Süel GM (2017) SnapShot: Electrochemical Communication in Biofilms. Cell 170(1):214–214.e1.

3. Prindle A, et al. (2015) Ion channels enable electrical communication in bacterial communities. Nature 527(7576):59–63.

4. Sirec T, Buffard P, Garcia-Ojalvo J, Asally M (2018) Electrical-charge accumulation enables integrative quality control during B. subtilis sporulation. bioRxiv:349654.

5. Piccolino M (1997) Luigi Galvani and animal electricity: Two centuries after the foundation of electrophysiology. Trends Neurosci 20(10):443–448.

6. McCaig CD, Rajnicek AM, Song B, Zhao M (2005) Controlling cell behavior electrically: current views and future potential. Physiol Rev 85(3):943–978.

7. Chang F, Minc N (2014) Electrochemical Control of Cell and Tissue Polarity. Annu Rev Cell Dev Biol 30(1):317–336.

8. Robinson KR (1985) The responses of cells to electrical fields: A review. J Cell Biol 101(6):2023–2027.

9. Adams DS, Levin M (2013) Endogenous voltage gradients as mediators of cell-cell communication: strategies for investigating bioelectrical signals during pattern formation. Cell Tissue Res 352(1):95–122.

10. Levin M (2014) Molecular bioelectricity: how endogenous voltage potentials control cell behavior and instruct pattern regulation in vivo. Mol Biol Cell 25(24):3835–3850.

11. Levin M (2007) Large-scale biophysics: ion flows and regeneration. Trends Cell Biol 17(6):261–270.

12. Lobikin M, Chernet B, Lobo D, Levin M (2012) Resting potential, oncogene-induced tumorigenesis, and metastasis: the bioelectric basis of cancer in vivo. Phys Biol 9(6):065002.

13. Yang M, Brackenbury WJ (2013) Membrane potential and cancer progression. Front Physiol 4:185.

14. Pardo LA, Stühmer W (2014) The roles of K + channels in cancer. Nat Rev Cancer 14(1):39–48.

15. Markx GH (2008) The use of electric fields in tissue engineering: A review. Organogenesis 4(1):11–17.

16. Tandon N, et al. (2009) Electrical stimulation systems for cardiac tissue engineering. Nat Protoc 4(2):155–173.

17. Levin M, Pezzulo G, Finkelstein JM (2017) Endogenous Bioelectric Signaling Networks: Exploiting Voltage Gradients for Control of Growth and Form. Annu Rev Biomed Eng 19(1):353–387.

18. Hunckler J, de Mel A (2017) A current affair: Electrotherapy in wound healing. J Multidiscip Healthc 10:179–194.

19. Famm K, Litt B, Tracey KJ, Boyden ES, Slaoui M (2013) A jump-start for electroceuticals. Nature 496(7444):159–161.

20. Hfilsheger H, Potel J, Niemann E-G (1981) Radiation and Environmental Biophysics Killing of Bacteria with Electric Pulses of High Field Strength. Radiat Env Biophys 20:53–65.

21. Chassy BM, Mercenier A, Flickinger J (1988) Transformation of bacteria by electroporation. Trends Biotechnol 6(12):303–309.

22. Schloss AC, et al. (2016) Fabrication of Modularly Functionalizable Microcapsules Using Protein-Based Technologies. ACS Biomater Sci Eng 2(11):1856–1861.

23. Marszalek P, Liu DS, Tsong TY (1990) Schwan equation and transmembrane potential induced by alternating electric field. Biophys J 58(4):1053–1058.

24. Kotnik T, Miklavcic D (2000) Analytical description of transmembrane voltage induced by electric fields on spheroidal cells. Biophys J 79(2):670–679.

25. Felle H, Porter JS, Slayman CL, Kaback HR (1980) Quantitative measurements of membrane potential in Escherichia coli. Biochemistry 19(15):3585–90.

26. Ramos S, Schuldiner S, Kaback HR (1976) The electrochemical gradient of protons and its relationship to active transport in Escherichia coli membrane vesicles. Proc Natl Acad Sci 73(6):1892–6.

27. Zheng J, Trudeau MC (2015) Handbook of Ion Channels eds Zheng J, Trudeau M (CRC Press) Available at: https://www.crcpress.com/Handbook-of-Ion-Channels/Zheng-Trudeau/9781466551404.

28. Kaim G, Dimroth P (1999) ATP synthesis by F-type ATP-synthase is obligatory dependent on the transmembrane voltage. EMBO J 18(15):4118–4127.

29. Strahl H, Hamoen LW (2010) Membrane potential is important for bacterial cell division. Proc Natl Acad Sci 107(27):12281–12286.

30. Milo R, Phillips R (2015) Cell Biology By the Numbers (Garland Science) Available at: http://book.bionumbers.org/.

31. Mason DJ, Lopéz-Amorós R, Allman R, Stark JM, Lloyd D (1995) The ability of membrane potential dyes and calcafluor white to distinguish between viable and non-viable bacteria. J Appl Bacteriol 78(3):309–15.

32. Breeuwer P, Abee T (2000) Assessment of viability of microorganisms employing fluorescence techniques. Int J Food Microbiol 55:193–200.

33. Cléach J, et al. (2018) Use of Ratiometric Probes with a Spectrofluorometer for Bacterial Viability Measurement. J Microbiol Biotechnol 28(11):1782–1790.

34. Kralj JM, Hochbaum DR, Douglass AD, Cohen AE (2011) Electrical spiking in Escherichia coli probed with a fluorescent voltage-indicating protein. Science 333(6040):345–8.

35. Alteri CJ, Lindner JR, Reiss DJ, Smith SN, Mobley HLT (2011) The broadly conserved regulator PhoP links pathogen virulence and membrane potential in Escherichia coli. Mol Microbiol 82(1):145–163.

36. Sträuber H, Müller S (2010) Viability states of bacteria--specific mechanisms of selected probes. Cytometry A 77(7):623–34.

37. Zhang Z, Milias-Argeitis A, Heinemann M (2018) Dynamic single-cell NAD(P)H measurement reveals oscillatory metabolism throughout the E. coli cell division cycle. Sci Rep 8(1):1–10.

38. Shinar G, Milo R, Martinez MR, Alon U (2007) Input output robustness in simple bacterial signaling systems. Proc Natl Acad Sci 104(50):19931–19935.

39. Kubitschek HE (1969) Growth during the bacterial cell cycle: analysis of cell size distribution. Biophys J 9(6):792–809.

40. Jayaram DT, Luo Q, Thourson SB, Finlay AH, Payne CK (2017) Controlling the Resting Membrane Potential of Cells with Conducting Polymer Microwires. Small 13(27):1700789.

41. Ren D (2001) A Prokaryotic Voltage-Gated Sodium Channel. Science 294(5550):2372–2375.

42. Payandeh J, Jr DLM (2015) Bacterial Voltage-Gated Sodium Channels (BacNa Vs) from the Soil, Sea, and Salt Lakes Enlighten Molecular Mechanisms of Electrical Signaling and Pharmacology in the Brain and Heart. J Mol Biol 427(1):3–30.

43. Murdoch LE, MacLean M, Endarko E, MacGregor SJ, Anderson JG (2012) Bactericidal effects of 405nm light exposure demonstrated by inactivation of escherichia, salmonella, Shigella, Listeria, and mycobacterium species in liquid suspensions and on exposed surfaces. Sci World J 2012. doi:10.1100/2012/137805.

44. MacLean M, Murdoch LE, MacGregor SJ, Anderson JG (2013) Sporicidal effects of high-intensity 405 nm visible light on endospore-forming bacteria. Photochem Photobiol 89(1):120–126.

45. FitzHugh R (1961) Impulses and Physiological States in Theoretical Models of Nerve Membrane. Biophys J 1(6):445–466.

46. Mallot HA (2013) Computational Neuroscience (Springer International Publishing, Heidelberg) doi:10.1007/978-3-319-00861-5.

47. Nikaido H (1989) Outer membrane barrier as a mechanism of resistance. Antimicrob Agents Chemother 33(11):1831–1836.

48. Verstraeten N, et al. (2015) Obg and Membrane Depolarization Are Part of a Microbial Bet-Hedging Strategy that Leads to Antibiotic Tolerance. Mol Cell 59(1):9–21.

49. Damper PD, Epstein W (1981) Role of the membrane potential in bacterial resistance to aminoglycoside antibiotics. Antimicrob Agents Chemother 20(6):803–8.

50. Allison KR, Brynildsen MP, Collins JJ (2011) Heterogeneous bacterial persisters and engineering approaches to eliminate them. Curr Opin Microbiol 14(5):593–598.

51. Allison KR, Brynildsen MP, Collins JJ (2011) Metabolite-enabled eradication of bacterial persisters by aminoglycosides. Nature 473(7346):216–220.

52. Atkinson JT, Campbell I, Bennett GN, Silberg JJ (2016) Cellular Assays for Ferredoxins: A Strategy for Understanding Electron Flow through Protein Carriers That Link Metabolic Pathways. Biochemistry 55(51):7047–7064.

53. Choi O, Kim T, Woo HM, Um Y (2015) Electricity-driven metabolic shift through direct electron uptake by electroactive heterotroph Clostridium pasteurianum. Sci Rep 4(1):6961.

54. Zerfaß C, Asally M, Soyer OS (2018) Interrogating metabolism as an electron flow system. Curr Opin Syst Biol 13:59–67.

55. Vieira-Pires RS, Szollosi A, Morais-Cabral JH (2013) The structure of the KtrAB potassium transporter. Nature 496(7445):323–328.

56. Haupt M, Bramkamp M, Coles M, Kessler H, Altendorf K (2006) Prokaryotic Kdp-ATPase: Recent insights into the structure and function of KdpB. J Mol Microbiol Biotechnol 10(2–4):120–131.

57. Ahmed A, Rushworth J V., Hirst NA, Millner PA (2014) Biosensors for whole-cell bacterial detection. Clin Microbiol Rev 27(3):631–646.

58. Asally M, et al. (2012) Localized cell death focuses mechanical forces during 3D patterning in a biofilm. Proc Natl Acad Sci 109(46):18891–6.

59. Young JW, et al. (2012) Measuring single-cell gene expression dynamics in bacteria using fluorescence time-lapse microscopy. Nat Protoc 7(1):80–8.

